# A human gut-BBB-brain microphysiological model for studying neurodegenerative diseases

**DOI:** 10.64898/2026.02.24.707716

**Authors:** Yu-Shan Deng, Wan-Peng Wang, Fei Wang, Guo-Ming Ma, Jian-Feng Lin, Chen-Xi Yan, Yi-Ling Zhou, Lu-Yao Wang, Xue-Qi Gong, Li Sun, Jian Zhao, Gang Pei, Lin Zhang, Wen-Yuan Wang

**Affiliations:** Interdisciplinary Research Center on Biology and Chemistry, Shanghai Institute of Organic Chemistry, Chinese Academy of Sciences, Shanghai, 200032, China; University of Chinese Academy of Sciences, Beijing, 100049, China; School of Life Sciences, Fudan University, Shanghai 200438, China; Shanghai Institute for Advanced Immunochemical Studies, ShangaiTech University, Shanghai, 201210, China; State Key Laboratory of Cell Biology, Shanghai Institute of Biochemistry and Cell Biology, Center for Excellence in Molecular Cell Science, Chinese Academy of Sciences, Shanghai 200031, China; The International Peace Maternity and Child Health Hospital, School of Medicine, Shanghai Jiao Tong University, Shanghai 200030, China

**Keywords:** Neurodegenerative diseases, gut-brain axis, organoids, microphysiological system

## Abstract

The gut-brain axis has emerged as a crucial factor in neurodegeneration, with growing evidence linking gut dysbiosis and metabolic dysfunction to Alzheimer’s disease (AD) progression. Unfortunately, the lack of human-relevant *in vitro* models limits our ability to effectively explore the mechanism of this axis. To address this gap, we have developed a human induced pluripotent stem cell (iPSC)-derived gut-blood-brain barrier (BBB)-brain microphysiological system that enables systematic investigation of gut-brain interaction in the context of AD under controlled conditions. Our findings reveal that the interaction between gut and brain organoids can promote the maturation of brain organoids, making them more similar to their physiological characteristics *in vivo*. Additionally, co-culture gut and brain organoids better recapitulates the pathological features of AD. We also discovered that gut organoids of AD can trigger neurodegenerative disease manifestations in healthy brain organoids. In summary, our microphysiological system provides a novel and versatile *in vitro* platform for studying the interaction between the gut and brain in neurodegenerative diseases.

## 1. Introduction

Neurodegenerative diseases (NDs), like Alzheimer’s disease, have traditionally been seen as disorders that mainly impact the central nervous system (CNS), characterized by progressive neuronal degeneration and functional decline^1^. However, recent researches reveal that NDs have a systemic nature, with growing evidence connecting peripheral abnormalities to CNS pathology^2^.

The gut-brain axis (GBA) is a complex bidirectional communication network that involves neural, endocrine, immune, and metabolic pathways^3^. Recent studies have demonstrated that gut microbiota dysbiosis is correlated with increased neuroinflammation and accelerated neurodegeneration^4^. Gut-derived metabolites have been shown to significantly enhance cognitive functions and alleviate AD-like pathology in the 5xFAD mouse model^5^. On the other hand, gut-derived pathological proteins, such as β-amyloid, may spread to the CNS through systemic circulation and neural pathways^6,7^.

Currently, studies investigating the roles of the gut-brain axis in neurodegenerative disorders have been predominantly based on longitudinal clinical studies and animal models^3^. While critical for understanding disease trajectories, clinical studies often struggle to keep pace with the needs of therapeutic development. Meanwhile, animal models often lack translational relevance due to differences between species^8^. The development of human *ex vivo* modeling, such as microphysiological systems (MPS), now enables the recapitulation of complex facets of human disease etiology under precisely controlled conditions^9^. By integrating engineered human tissues, fluidic channels, and multi-organ interactions, MPS enables (1) physiologically relevant modeling of bidirectional gut-brain crosstalk through neuronal, endocrine, and immune pathways; (2) high-precision manipulation of microbiome metabolites, barrier integrity, and neuroactive compounds in a human-relevant context; and (3) superior capabilities for drug testing, allowing high-throughput screening of compound efficacy and toxicity in a human-relevant system while reducing reliance on animal models^10–12^. Notably, integrated platforms that combine gut, blood-brain barrier, and neural components have successfully modeled PD pathology, highlighting the potential of these approaches for studying neurodegenerative diseases^13^. The PEGASO multi-organ platform enables reliable *in vitro* simulations of donepezil pharmacokinetics and efficacy, demonstrating significant promise for preclinical research and optimizing drug development^14^. While capable of mimicking certain human physiological features, these models remain constrained in their ability to reproduce phenotype because they rely on 2D culture systems that lack the physiological complexity of *in vivo* tissues. Compared with 2D cell culture, 3D organoids more accurately mimic *in vivo* tissue architectures, cellular interactions, and disease pathophysiology, offering superior physiological relevance for biomedical research^15–17^.

To overcome current limitations and improve our understanding of the gut-brain axis involvement in neurodegeneration, we have developed an advanced *in vitro* platform that integrates human induced pluripotent stem cell (iPSC)-derived gut, blood-brain barrier, and brain organoids within a circulatory system, enabling physiological communication. Our platform enhances organoid maturation through tissue interactions, successfully recapitulating disease-specific pathologies by facilitating gut-brain communication. This innovative system serves as a powerful tool for investigating neurodegenerative mechanisms and developing therapeutic strategies, providing new insights into the role of the gut-brain axis in disease progression.

## 2. Results

### 2.1. Construction of the human gut-BBB-brain model

From a physiological perspective, we simplified the gut-brain axis (GBA) into three main components: the gut, the blood-brain barrier, and the brain (**Figure 1A**). The human gut and brain are complex organs with a variety of cell types and unique structures. Human organoids can recreate the architecture and physiology of human organs, providing unique opportunities to study human diseases and complementing existing model systems^18^. Thus, we decided to differentiate human iPSCs into gut and brain organoids to better recapitulate the *in vivo* physiological state of these two organs. The blood-brain barrier plays a key role in maintaining brain homeostasis by controlling the entry of harmful molecules into the brain^19^.

**Figure 1.**
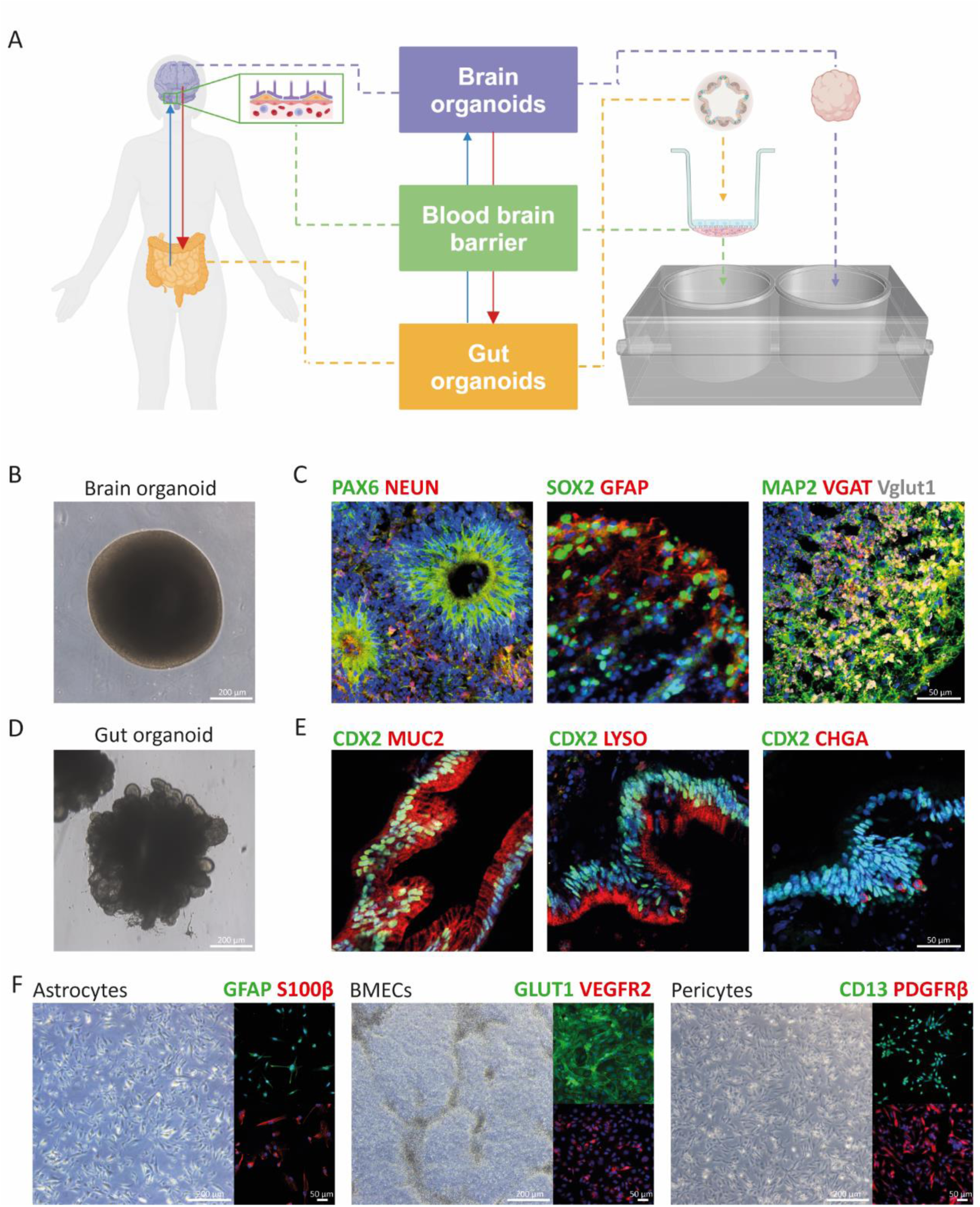
Human GBB microphysiological model. (A) Schematic representation of model design. (B) Bright field image of a brain organoid. Scale bar, 200 μm. (C) Cryosections of HCSs stained with the indicated markers. Scale bar, 50 μm.(D) Bright field image of a gut organoid. Scale bar, 200 μm. (E) Cryosections of HIOs stained with the indicated markers. LYSO: lysozyme; CHGA: chromogranin A. Scale bar, 50 μm. (F) Left image: induced astrocytes marked with GFAP and S100β. Middle image: induced BMECs marked with GLUT1 and VEGFR2. Right image: induced pericytes marked with CD13 and PDGFRβ. Scale bar, 200 μm for bright field images; 50 μm for immunofluorescence images.

The BBB is composed of brain microvascular endothelium cells that line the cerebral capillaries, surrounded by pericytes and astrocyte end feet^20^. Therefore, we seeded iPSC-derived brain microvascular endothelium cells (BMECs), astrocytes (ACs), and pericytes (PCs) in a specific pattern in 12-well trans-well plates. All organoids and the BBB were matured individually before integration to establish specific features and physiological functions. The co-culture chip is manufactured from acrylic and is biocompatible with human tissue. The culture chamber has open access and can be connected in any order to model different physiological scenarios. The microphysiological platform uses a multi-channel peristaltic pump with six individual channels, enabling up to six chips to be used simultaneously. The chip design enables the BBB to be constructed on a trans-well and placed into the culture chamber after it has passed quality control, allowing it to perform its normal barrier functions.

Brain organoids, also referred to as human cortical spheroids (HCSs), exhibited a diameter of up to 2 mm after 100 days of induction, with smooth edges and an opaque, milky-white appearance (Figure 1B). Immunofluorescence analysis confirmed the presence of neurons (NeuN+ and MAP2+), astrocytes (GFAP+), and neural stem cells (PAX6+ and SOX2+) in HCSs (Figure 1C). Additionally, markers for excitatory neurons (Vglut+) and inhibitory neurons (VGAT+) were successfully expressed in the organoids (Figure 1C). Notably, PAX6+ neural stem cells aggregated into rosette-like structures, recapitulating the early morphology of the neural tube during embryonic development, which serves as a critical indicator of the organoids’ developmental potential (Figure 1C). These staining results demonstrated that the iPSC-derived HCSs possess a complex cellular composition and recapitulate key features of brain development.

In parallel, induced human intestinal organoids (HIOs) displayed prominent lumen-like structures during induction, morphologically resembling intestinal tissues (Figure 1D). Immunofluorescence revealed that CDX2, a marker of intestinal epithelial cells, was exclusively expressed along the luminal wall (Figure 1E). Furthermore, the organoids secreted muc2 (a goblet cell marker), lysozyme (a Paneth cell marker), and chromogranin A (an enteroendocrine cell marker) (Figure 1E), indicating that the iPSC-derived intestinal organoids not only mimic the structural organization of *in vivo* intestines but also exhibit functional secretory properties.

Subsequently, iPSCs were differentiated into astrocytes, BMECs, and pericytes, followed by immunofluorescence validation of their respective biomarkers. Astrocytes, cultured in 2D conditions, were positive for GFAP and S100β (Figure 1F). BMECs formed tightly interconnected monolayers of flat cells and expressed endothelial markers VEGFR2 and Glut1 (Figure 1F). Pericytes exhibited spindle-shaped somas and expressed CD13 and PDGFRβ (Figure 1F). These results confirm the successful derivation of astrocytes, BMECs, and pericytes from iPSCs, laying the cellular foundation for constructing an iPSC-derived blood-brain barrier model.

### 2.2. Generating a functional blood-brain barrier

To recreate a human BBB *in vitro*, we seeded BMECs on the apical side of a trans-well to represent the “blood side”. On the basal side, we embedded ACs and PCs in Matrigel at a 1:1 ratio to form a human iPSC-derived BBB (iBBB) (**Figure 2A**). Trans-endothelial electrical resistance (TEER) serves as a reliable and intuitive indicator of BBB integrity. We measured the TEER value of the iBBB during the culture process using an EVOM2 device. Notably, the TEER values of our iBBB reached as high as 2000 Ω·cm^2^, maintaining levels above 1000 Ω·cm^2^ for about 30 days. This demonstrated that the iBBB achieved physiologically relevant values^21^(Figure 2B). Additionally, we observed clear expression of specific markers for ACs (S100β+), PCs (TAGLN+), and BMECs (VEGFR2+), indicating that the distinct characteristics of the different cell types were still preserved even after long-term co-culture (Figure 2C). Furthermore, the tight junction protein ZO-1 was localized at the cell boundaries, demonstrating the formation of tight cell connections, which are crucial for BBB integrity^22^ (Figure 2C).

**Figure 2.**
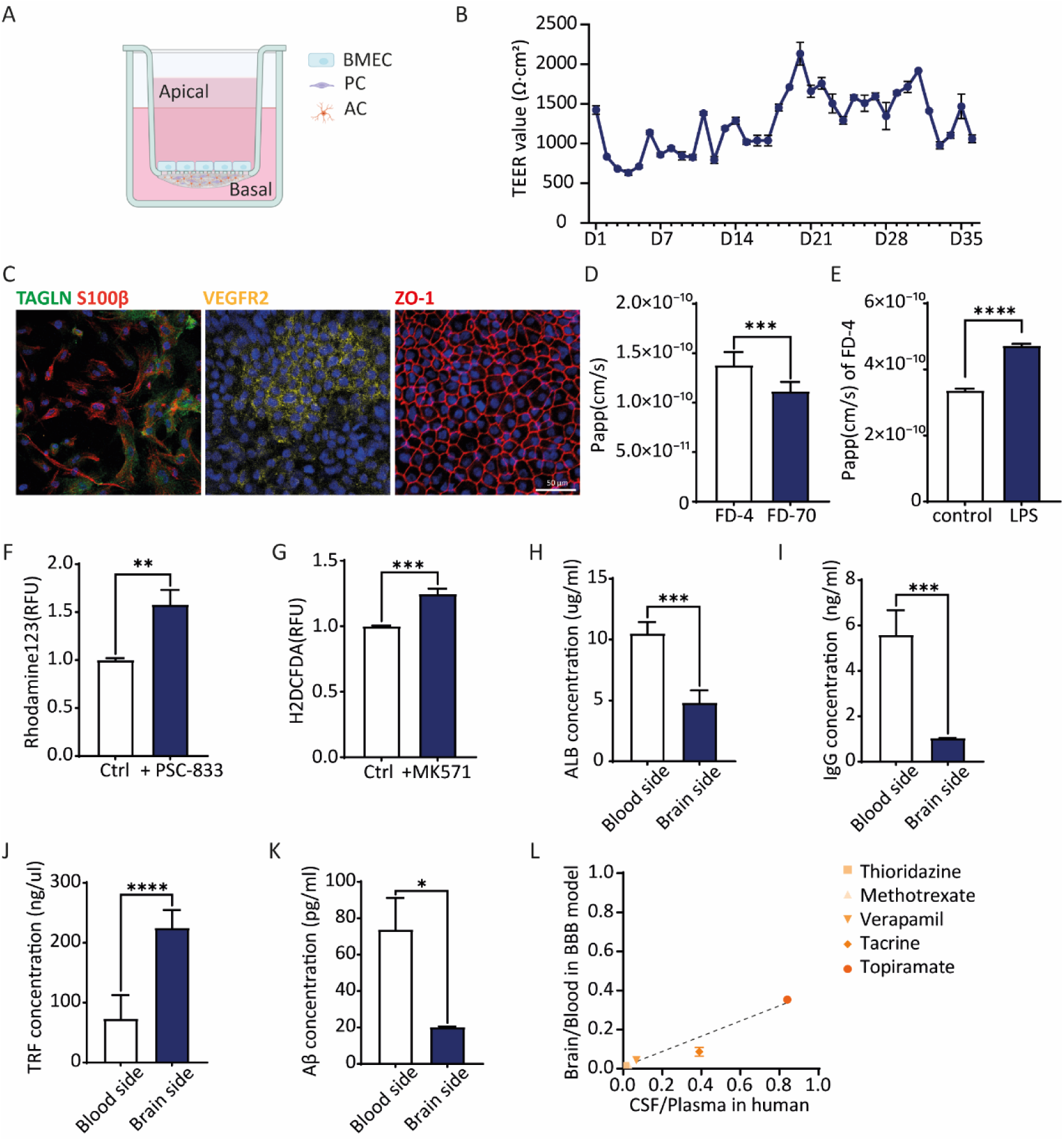
Identification of physiological properties and functions of the human iBBB. (A) Schematic of the iBBB seeding strategy. BMECs are seeded on the apical side of the trans-well; ACs and PCs are embedded in Matrigel and seeded on the basal side of the trans-well. (B) TEER measurement of the iBBB lasts for 35 days. (C) Immunofluorescence images of PCs (TAGLN+), ACs (S100β+), and BMECs (VEGFR2+, ZO1+) in the iBBB model. Scale bar, 50 μm. (D) iBBB permeability coefficients of FD-4 and FD-70. Papp: apparent permeability. (E) iBBB permeability coefficients of FD-4 with or without the treatment of LPS. (F and G) Efflux pump function measured through rhodamine123 as a substrate of P-gp and H2DCFDA as a substrate of MRPs, with or without the addition of efflux transporter inhibitors, which PSC-833 for P-gp and MK571 for MRPs. RFU: relative fluorescence unit. (H-K) Apical-to-basal filtration of the human albumin (ALB, 40 ug/ml), immunoglobulin (IgG; 10 ug/ml), transferrin (TRF;2.5 ug/ml), and beta-amyloid (Aβ; 1ug/ml). (L) Drug permeability tests of the iBBB model versus *in vivo* human drug permeability (y= 0.3911x - 0.002654, R^2^=0.9198).

To assess the permeability of the iBBB, we treated it with the proteins of different molecular sizes, specifically 4 kDa and 70 kDa fluorescein isothiocyanate (FITC) labeled dextran (FD-4 and FD-70, respectively). The results showed that the iBBB model exhibited higher permeability to FD-4 compared to FD-70 (Figure 2D), consistent with the general understanding that permeability decreases with increasing molecular weight^23^. Additionally, treatment with LPS significantly increased the blood-to-brain leakage of FD-4 compared to conditions without LPS (Figure 2E), suggesting that our iBBB model can respond to pro-inflammatory substances and exhibits an impaired barrier phenotype.

The activities of efflux pump proteins, such as P-glycoprotein (P-gp) and multidrug resistance proteins (MRPs), are the key features of the BBB *in vivo*. These proteins can recognize various substrates and pump them back into the bloodstream, thereby playing an important role in protecting brain tissue from exogenous compounds, including many drugs^19^. To verify the function of efflux pumps in our *in vitro* iBBB models, we pretreated the iBBB with the inhibitors PSC-833 and MK571, which target P-gp and MRPs, respectively. We then monitored the penetration of specific substrates, Rhodamine123 and 2’,7’-Dichlorodihydrofluorescein diacetate (H2DCFDA), across the iBBB. The treatment with inhibitors significantly increased the transport of these substrates from the apical to the basal side (Figure 2F, G), indicating that P-gp and MPRs were functionally active in the iBBB model.

Among the most abundant proteins in human blood are Albumin (ALB), immunoglobulin (IgG), and transferrin (TRF). While ALB and IgG are primarily restricted to the blood side of the barrier, TRF can cross the BBB through receptor-mediated transcytosis^24^. To further validate the physiological functions of the iBBB, we investigated whether these molecules could be selectively filtered. As expected, ALB and IgG remained confined to the blood side, while TRF accumulated in the brain side (Figure 2H-J). These results suggest that our iBBB model can replicate receptor-mediated transcytosis and selectively transport different molecules.

Additionally, we measured the permeability of the iBBB to amyloid beta (Aβ), a pathological protein associated with Alzheimer’s disease that can originate from peripheral organs, particularly the gut^6^. We found that more Aβ was detected in the blood side (Figure 2K), indicating that the iBBB model can restrict Aβ entry into the brain.

The blood-brain barrier serves to protect the central nervous system while also limiting the delivery of most drugs^25^. We investigated the correlation between *in vivo* human drug permeability and drug permeability in the *in vitro* iBBB model. The drug permeability observed in the *in vitro* BBB model was largely consistent with *in vivo* data^26^, demonstrating a strong correlation (R^2^= 0.9198) (Figure 2L).

In summary, these results demonstrate that our iBBB model reproduces functions similar to those found *in vivo*, including selective permeability, active efflux pumps, and receptor-mediated transcytosis. Therefore, it is a suitable tool for predicting drug permeability.

### 2.3. Gut-BBB-brain co-culture HCSs exhibit a higher physiological correlation with the *in vivo* human brain

To investigate the physiological consequences of communication among the gut, BBB, and brain, we established three distinct culture systems: (1) isolated HCSs (brain organoids only, B), (2) a co-culture of HIOs and HCSs (gut-brain, GB) representing the direct gut-brain interaction, and (3) a co-culture of HIOs, iBBB, and HCSs (gut-BBB-brain, GBB) (**Figure 3A**). After four days of co-culture, we collected the HCSs for further analysis. Three healthy iPSC lines were selected to generate these systems, which we abbreviated as HC_B, HC_GB, and HC_GBB, respectively.

**Figure 3.**
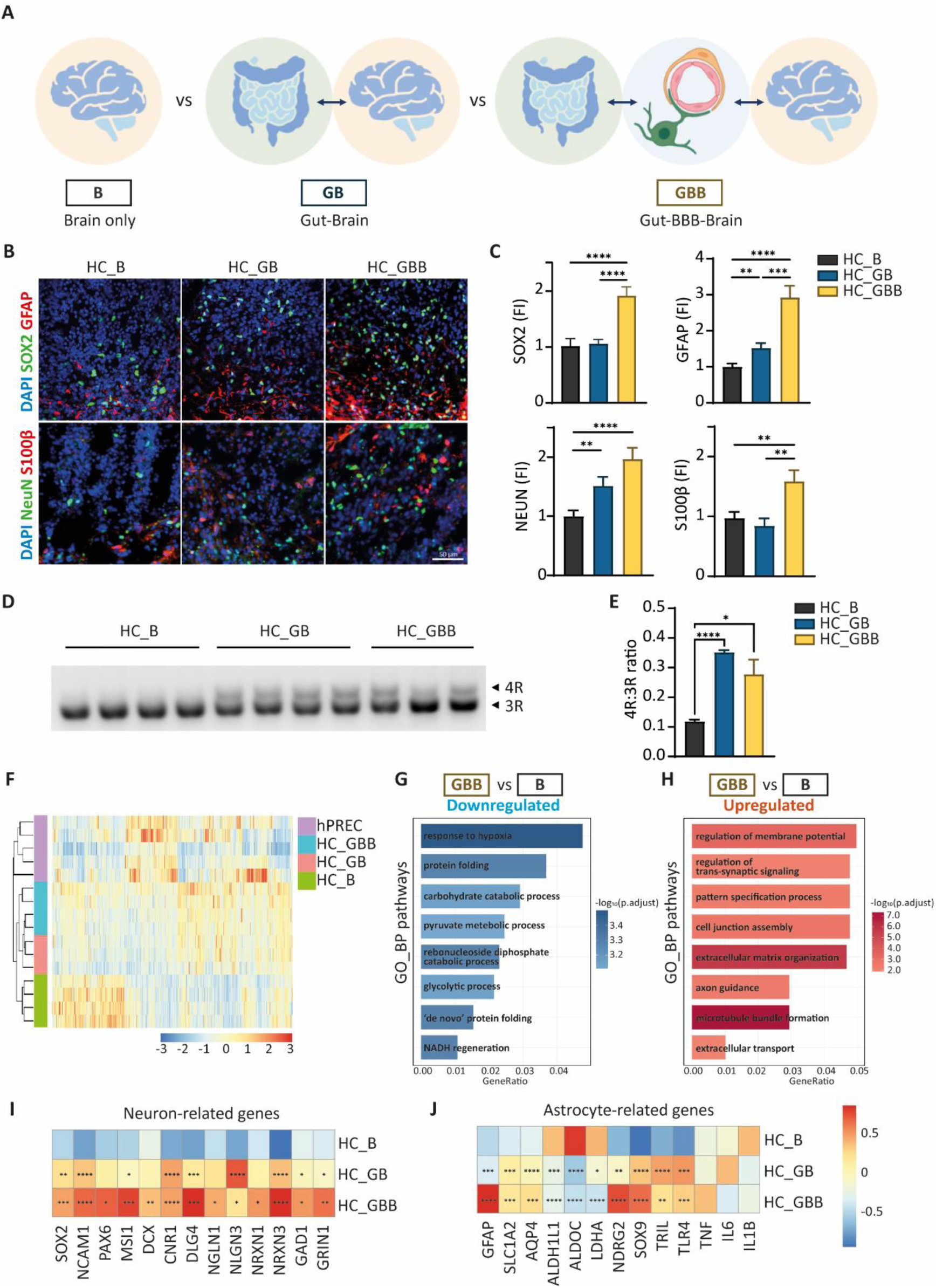
GBB-culture changes HCSs’ transcriptional features, developmental stages, and maturity of cells. (A) Schematic presentation of different culture models. (B) Cryosections of HCSs in different culture models stained with SOX2, GFAP, NEUN, and S100β. Scale bar, 50 μm. (C) Fluorescence intensity of SOX2, GFAP, NeuN, and S100β. FI: fluorescence intensity. (D) Semi-quantitative PCR of 3R and 4R tau expression in HC_HCSs of different culture models. (E) Quantification of the 4R to 3R ratio of HCSs grouped by HC_B, HC_GB, and HC_GBB. (F) Heatmap of transcriptional analysis of HCSs in different culture modes and an adult human dataset. (G and H) GO pathway enrichment in the HC_GBB HCSs over those in the HC_B group. (I and J) DEGs of (I) neuron-related and (J) astrocyte-related genes among HC_B, HC_GB, and HC_GBB. Analysis was carried out between HC_GB versus HC_B and HC_GBB versus HC_B.

We analyzed cellular markers using immunofluorescence staining. The results indicated a significant increase in NeuN-positive neurons in the HC_GB and HC_GBB groups compared to the HC_B group. In addition, the neuronal stem cell marker SOX2 and the astrocyte markers GFAP and S100β were significantly elevated in the GBB condition compared to the other two groups (Figure 3B, C). These findings suggest that the interaction among the gut, BBB, and brain significantly impacts the proliferation and maturation of neural stem cells and astrocytes.

As reported, iPSC-derived HCSs typically resemble the mid-late stages of fetal cortical development, which limits their utility for investigating the mechanisms underlying neurodegenerative diseases^27,28^. Alternative splicing of the tau protein generates two isoforms that differ due to the splicing of exon 10, which is responsible for encoding the second MT-binding repeat, leading to the generation of either three-repeat (3R) or four-repeat (4R) tau^29^. Only 3R tau is expressed during the fetal and early postnatal period, while the expression of 4R tau increases with development^29,30^. In this study, we used the 4R to 3R ratio as an indicator of the developmental stage. Semi-quantitative PCR analysis showed that the 4R/3R tau ratio approached unity in the HC_GB (0.35 ± 0.02) and HC_GBB (0.28 ± 0.09) groups, which was significantly higher than in the HC_B (0.12 ± 0.01), suggesting that co-culture with HIOs promotes the development of HCSs (Figure 3D, E).

Furthermore, transcriptome sequencing revealed that the interaction among the gut, BBB, and brain markedly affected the transcriptional features of HCSs. This was evident from gene clustering and principal component analysis (PCA) analysis (Figure 3F and Figure S1A). Moreover, to determine which model is more likely to restore the *in vivo* physiological state, we compared our data with an adult human brain sequencing dataset (PRJNA720779^31^). The results indicated that GBB-cultured HCSs exhibited transcriptional profiles more similar to those of the human dataset, as shown in the sample distance analysis (Figure 3F and Figure S1B).

Compared with the HC_B group, we identified 4580 and 4972 differentially expressed genes (DEGs) in the HC_GB and HC_GBB groups, respectively (Figure S1C). However, few DEGs were identified between the HC_GB and HC_GBB groups, which aligned with the clustering results (Figure 3F and Figure S1C). Subsequent analysis of DEGs using GO pathway enrichment showed that many metabolism-related pathways, such as carbohydrate catabolism, pyruvate metabolism, and glycolysis, were downregulated in the GBB-culture group compared to isolated HCSs (Figure 3G). Additionally, pathways related to hypoxia response and protein folding were also decreased in the HC_GBB group (Figure 3G). Conversely, upregulated genes were mainly enriched in pathways related to synaptic functions, such as membrane potential regulation, synaptic signaling, axon guidance, and intercellular communication, suggesting enhanced neural activity and communication (Figure 3H).

When comparing the HC_GB and HC_B groups, protein folding-related pathways were significantly downregulated, while genes related to cell division were significantly upregulated (Figure S1D). Unexpectedly, despite the absence of immune cells in the iPSC-derived HCSs, the upregulated genes in the GBB group compared to the HC_GB group were enriched in many immune pathways (Figure S1E). Further analysis of these genes revealed that they play important roles in neuroinflammatory regulation and immune responses of astrocytes^32–38^, suggesting that the presence of the BBB may alter the immune environment of HCSs and influence the physiological state of astrocytes.

Moreover, DEGs analysis of neuron-related genes revealed elevated expression of genes critical for neurodevelopment (SOX2, NCAM1, PAX6, MSI1, DCX^39–42^) and synaptic function (CNR1, DLG4, NLGN1, NLGN3, NRNX1, NRNX3, GAD1, GRIN1^43–47^) in both the HC_GB and HC_GBB groups (Figure 3I). A marked increase in GFAP expression was observed in the HC_GBB group, while the expression of pro-inflammatory cytokines (TNF, IL6, IL1B^48^) did not change significantly, suggesting that GBB-culture could promote the proliferation and maturation of astrocytes in HCSs rather than inducing an inflammatory state (Figure 3J). Additionally, genes linked to energy metabolism and immune function in astrocytes were also affected^49–52^ (Figure 3J).

Previous transcriptomic analysis revealed that the addition of HIOs significantly influences metabolic pathways. Consequently, we performed metabolomic profiling of HCSs under different culture conditions. The GBB-culture enhanced the levels of amino acid-related metabolites in HCSs (Figure S2A). Integrating this finding with transcriptomic data, we hypothesize that GBB-culture increases amino acid levels in cerebral organoids, thereby promoting neurotransmitter synthesis. Interestingly, we observed that co-culture with HIOs initially increased the levels of multiple metabolites in HCSs. However, subsequent introduction of the iBBB restored these metabolites to baseline levels found in isolated HCSs, underscoring the vital role of the BBB in peripheral-CNS interactions (Figure S2B).

Overall, GBB-culture enhances the proliferation and differentiation of neurons and astrocytes, recapitulating adult-like transcriptional patterns and developmental stages. These findings collectively establish the GBB-culture system as a superior *in vitro* model for neurodegenerative diseas1e research, overcoming significant limitations of conventional organoid models by achieving adult-like maturation states while incorporating physiologically relevant inter-tissue interactions.

### 2.4. The GBB-culture model can recapitulate AD pathologies in HCSs better

To investigate how different culture systems influence the modeling of neurodegenerative diseases, we compared the phenotypes of Alzheimer’s disease and healthy controls across the three culture conditions. We utilized iPSCs derived from AD patients to establish different culture models^53–55^, which we abbreviated as AD_B, AD_GB, and AD_GBB, respectively, and compared these with healthy controls (**Figure 4A**).

**Figure 4.**
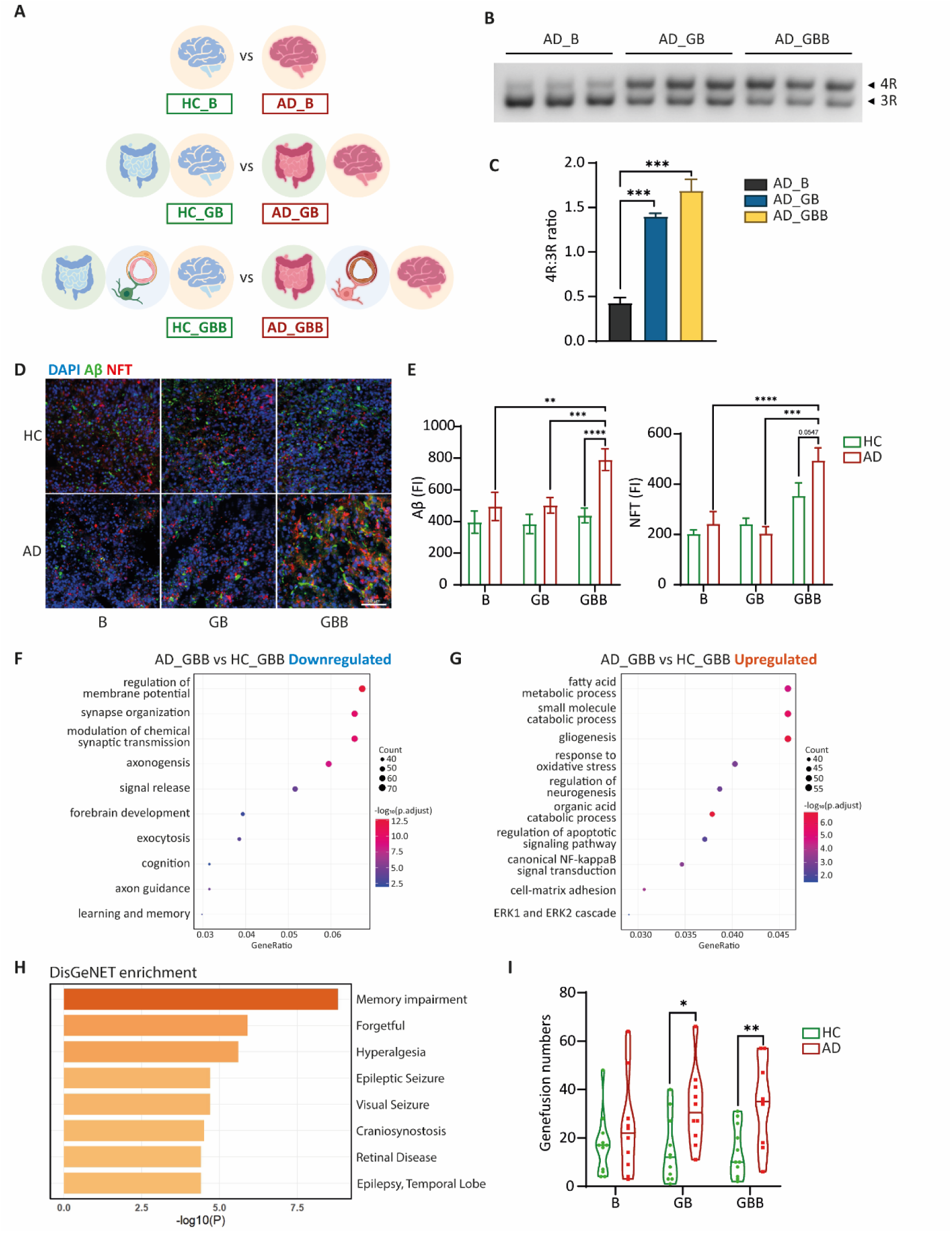
GBB co-culture increases pathology-related features in AD HCSs. (A) Schematic presentation of different conditions compared to AD with HC. (B) Semi-quantitative PCR of 3R and 4R tau expression in AD_HCSs of different culture models. (C) Quantification of the 4R to 3R ratio of HCSs grouped by AD_B, AD_GB, and AD_GBB. (D) Cryosections of HCSs in different culture models stained with Aβ and NFT. Scale bar, 50 μm. (E) Fluorescence intensity of Aβ and NFT in (D). (F and G) GO pathway enrichment of (F) downregulated genes and (G) upregulated genes in the AD_GBB HCSs over those in the HC_GBB group. (H) Disease correlation analysis of specific downregulated genes between AD and HC in the GBB system. (I) The number of fusion genes of HCSs in different culture models with AD and HC conditions.

In a healthy brain, the isoforms of tau protein, 3R tau and 4R tau, are present in approximately equal amounts. However, a higher ratio of 4R to 3R has been reported in AD, leading to pathological tau aggregation and the formation of neurofibrillary tangles (NFT) ^56,57^. When we compared the 4R to 3R ratio in the AD HCSs of the different culture models, we found that the ratio in AD_B was less than 0.5 (0.43 ± 0.10). In contrast, the ratios in the AD_GB and AD_GBB groups (1.40 ± 0.06 and 1.69 ± 0.20, respectively) were closer to 1.5 (Figure 4B, C), which is comparable to the ratios found in AD patients^58^.

Immunofluorescence analysis of AD pathological proteins revealed that only the GBB-culture system demonstrated a significant upregulation of both Aβ plaques and NFTs in AD-derived HCSs (Figure 4D, E). Additionally, the levels of these two pathological proteins were significantly higher in AD_GBB HCSs than in the AD_B and AD_GB groups, which were also supported by ELISA and western blot results (Figure S3A, B). These results validate that HCSs co-cultured with HIOs and BBB better recapitulate AD pathologies.

A comparison of DEGs between AD and HC in the various culture models revealed significant transcriptional differences (Figure S4A). We detected 3,423 downregulated genes between AD_B and HC_B (Figure S4A), although these genes were not significantly enriched in any specific pathway. Pathways related to axonogenesis, regulation of membrane potential, and trans-synaptic signaling, which were crucial for neural functions, were downregulated in the AD_GB and AD_GBB groups, indicating impaired neuronal function in AD (Figure 4F and Figure S4C). Moreover, there was a significant enrichment of downregulated pathways related to learning and memory, directly associated with the AD phenotype, particularly in the GBB group (p.adj = 0.004) compared to the GB group (p.adj = 0.038) (Figure 4F and Figure S4C). Conversely, pathways associated with gliogenesis were more highly expressed in all three culture models, correlating with higher levels of neuroinflammation in AD (Figure 4G and Figure S4B, D). Notably, the downregulation of cognition pathways and upregulation of ERK1 and ERK2 cascade pathways in AD HCSs were observed specifically within the GBB-culture system (Figure 4F, G). The upregulation of mitogen-activated protein kinases ERK1/2 is linked to the progression of neurofibrillary degeneration in Alzheimer’s disease^59^.

We conducted an intersection analysis of the differentially expressed genes detected between AD and HC in the different culture groups, identifying 425 genes that were specifically downregulated in the GBB group (Figure S4E). When we explored transcriptomic changes associated with disease patterns using the DisGeNET platform, we found that the genes specifically downregulated in AD_GBB were enriched in memory impairment and forgetfulness (Figure 4H).

Furthermore, we isolated the DEGs detected in AD HCSs compared to the healthy group only under GBB co-culture conditions, which included 425 downregulated genes and 336 upregulated genes. Notably, many of these genes have been previously reported to be associated with AD risk or pathology (ALOX5, APOA1, ALOX12, ATOH8, AZIN2, among others, see Table S1). Additionally, a significant number of solute carrier (SLC) family genes were identified among the DEGs (SLC9A5, SLC6A7, SLC4A9, SLC4A7, etc.), which are known to play crucial roles in AD pathogenesis^60^. We also discovered several genes with critical functions in the central nervous system that have not been thoroughly investigated in the context of AD (RASD1, RNF141, SLC35F6, SLC7A11, VWC2L, RBFOX3, NECAB2, MAGEL2, KCNS2, ITGAM, GRIN2D, GDPD5, GABRE, EHBP1L1, and CHRNA7), indicating their potential involvement in the disease.

Moreover, gene fusion events have been shown to increase in AD patients due to genomic instability^61^. As expected, the number of gene fusions was significantly higher in the AD group than in the HC group after the addition of HIOs co-culture (Figure 4I and Figure S4F).

Our data indicate that AD HCSs in the GBB system exhibit both pathological and transcriptional differences that more closely resemble those of AD. The GBB microphysiological platform is therefore a more suitable tool for systematically studying AD.

### 2.5. AD patient-derived HIOs can impair the physiological function of the healthy brain

To investigate the biological impact of AD-associated gut metabolites on the healthy brain, we replaced the HIOs in HC_GBB with AD patient-derived HIOs, creating a new model referred to as AD_G_HC_BB. We used HC_GBB as the control **(Figure 5A**).

**Figure 5.**
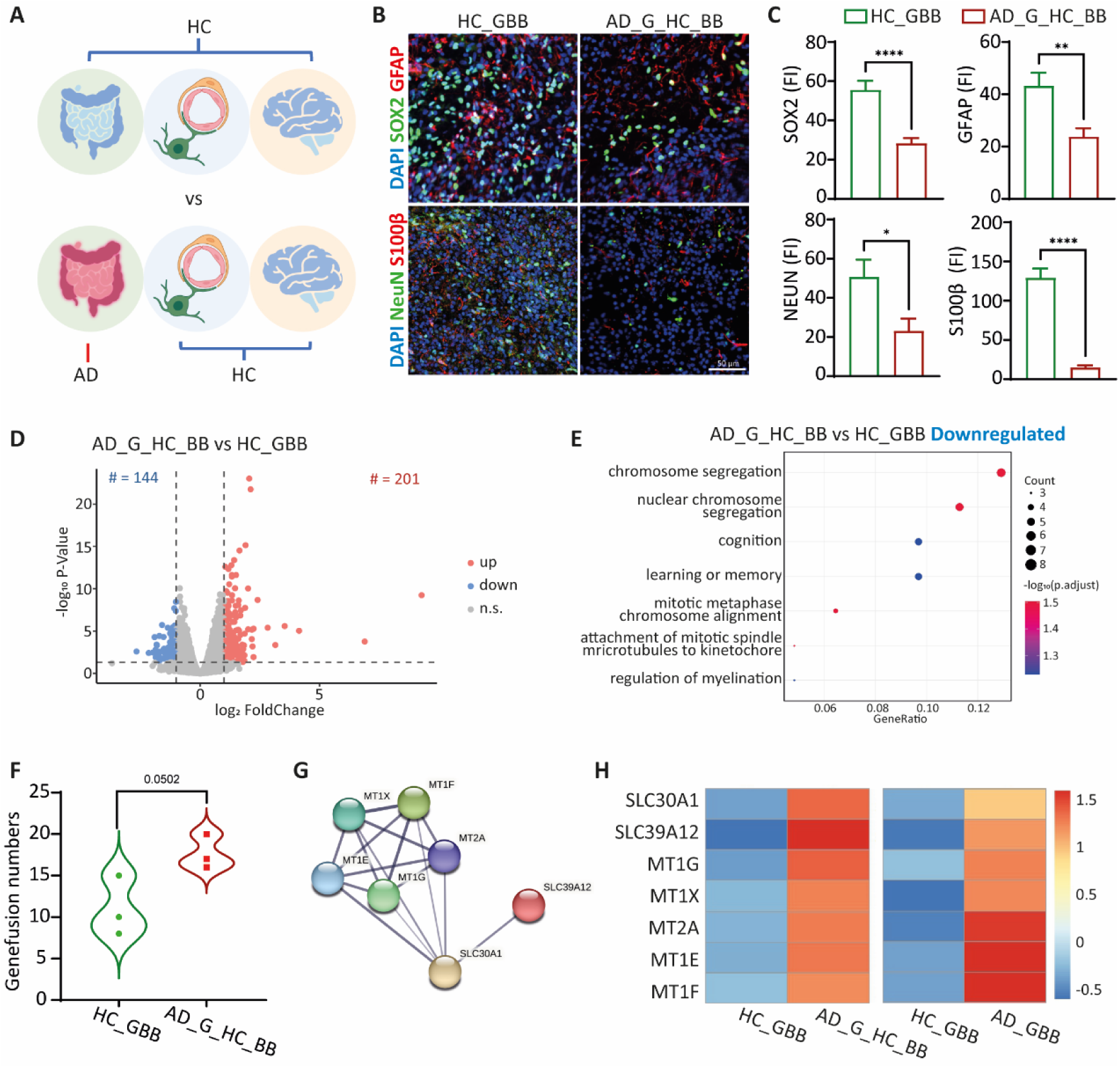
AD HIOs induce an AD-like pathological phenotype in HC HCS after co-culture. (A) Schematic presentation of AD HIOs co-culture with HC BBB and HC HCSs. (B) Cryosections of HCSs in HC_GBB and AD_G_HC_BB systems stained with SOX2, GFAP, NEUN, and S100β. Scale bar, 50 μm. (C) Fluorescence intensity of SOX2, GFAP, NEUN, and S100β in (B). (D) Volcano plots showing differential expression genes between AD_G_HC_BB HCSs and HC_GBB HCSs. (E) Downregulated GO pathway enrichment in the AD_G_HC_BB HCSs over those in the HC_GBB group. (F) The number of fusion genes of HCSs in the AD_G_HC_BB and HC_GBB groups. (G) PPI network for upregulated DEGs in AD_G_HC_BB HCSs. (H) Relative expression level of genes shown under (G) in AD_GBB and HC_GBB HCSs.

Levels of markers for neurons, neural stem cells, and astrocytes in HC HCSs were decreased significantly (Figure 5B, C). However, the levels of Aβ deposits and phosphorylated tau remained unchanged (Figure S5A, B). We hypothesized that 4 days of co-culture may not be sufficient to induce the deposition of pathological proteins typically associated with AD.

Additionally, AD HIOs induced notable changes in the transcriptome of HC HCSs. We identified a total of 345 differentially expressed genes between the two groups (Figure 5D). GO analysis revealed that the downregulated genes in the AD_G_HC_BB group were significantly enriched in pathways related to the cell cycle, which is consistent with the observed decrease in neural stem cells and astrocytes from the immunofluorescence analysis (Figure 5B, C, and E). Importantly, pathways associated with learning, memory, and cognition were also found to be markedly suppressed (Figure 5E), indicating that AD HIOs can elicit transcriptional changes in HC HCSs that resemble those found in AD. Moreover, co-culture with AD HIOs showed a trend towards an increase in the number of gene fusion events occurring in HC HCSs (Figure 5F), further reflecting the pathological characteristics associated with AD.

Subsequently, we performed an intersection analysis between the DEGs of the AD_GBB and HC_GBB groups, as well as those of the AD_G_HC_BB and HC_GBB groups. The results revealed that the common gene set included several metallothioneins (MT1X, MT1E, MT1F, MT1G, MT2A) and was enriched in pathways associated with metal homeostasis, which is disrupted in AD(Figure 5G)^62^. Additionally, the expression of these genes was significantly upregulated in both the AD_GBB and AD_G_HC_BB HCSs compared to the HC_GBB HCSs (Figure 5H). This finding is consistent with the expression pattern observed in AD patients^63^. We speculate that the gut may potentially influence AD pathology by modulating metal homeostasis in the brain under AD conditions.

Considering that in our system, the intestinal and brain organoids can only communicate via metabolic pathways, we conducted metabolome profiling analyses on AD and healthy HIOs. A total of 58 differential metabolites were detected between AD and HC HIOs, with 28 being upregulated and 30 downregulated (Figure S6A, B). The majority of these differential metabolites were related to benzene and its derivatives, amino acids and their metabolites, as well as glycerophospholipid (GP), which collectively accounted for half of the total (Figure S6C).

We identified several AD-related metabolites through disease correlation analysis: L-valine, L-isoleucine, Ornithine, hypoxanthine, and dihydroxyacetone phosphate (DHAP). Our data indicated that the expression levels of all five metabolites were markedly decreased in the AD HIOs (Figure S6D), which was consistent with existing literature on AD patients^64–67^, demonstrating that AD HIOs can effectively replicate the metabolic characteristics associated with AD *in vitro*.

In summary, our results indicate that AD HIOs can induce phenotypes similar to those seen in AD pathologies in HC HCSs through metabolic pathways, likely by affecting metal homeostasis. Our GBB microphysiological system could serve as a practical tool for studying the metabolic pathways involved in the gut-brain axis.

## 3. Discussion

Neurodegenerative disease studies are a global challenge in an aging society^68^. While most of our current knowledge about NDs mainly comes from post-mortem brain samples and longitudinal clinical data, supplemented with animal models and monolayer cell cultures, the complexity of neurodegenerative diseases demands innovative approaches that can address both cellular-level pathologies and systemic interactions^69^.

Recent advancements in organoid and multi-organ-on-a-chip technologies have come together to create advanced *organoid-on-a-chip* platforms. These innovations are transforming the study of complex NDs, such as Alzheimer’s disease^17^. These platforms overcome the significant limitations of traditional models by combining the cellular complexity of organoids with the integration of multiple organs. This combination enables the recreation of human organ-level physiology in both healthy and pathological states *in vitro*^17^.

Our study investigates the interaction between the gut and brain, both in healthy and pathological states. We aim to reconstruct their interactions *in vitro* to examine their effects on neuropathology under controlled conditions. To achieve this, we have developed a co-culture system that integrates gut organoids, a functional BBB, and brain organoids. This design closely resembles the physiological gut-brain axis, with the BBB acting as a selective gatekeeper - a feature often lacking in most existing models^70^.

The GBB model presents three significant advantages that greatly enhance current research capabilities: (1) improved physiological relevance: compared with existing gut-brain axis models^13,14^, we utilize organoids with higher complexity and add the BBB with excellent functions so that our model can accurately recapitulate essential *in vivo* multi-organ interactions, offering data that is more clinically predictive than conventional models; (2) exceptional modular flexibility: researchers can easily customize the platform for a variety of applications – ranging from disease modeling to drug development - by exchanging organoid types or using iPSCs from different donors; and (3) strong potential for personalized medicine: the ability to incorporate patient-specific iPSCs allows for the creation of individualized therapeutic screening platforms, which is a crucial for advancing precision medicine approaches in neurodegenerative disease research.

Our research demonstrates that the interactions among brain organoids, gut organoids, and the blood-brain barrier increase the expression of neurons, neural stem cells, and astrocyte markers, suggesting that the GBB-culture system promotes the proliferation and maturation of neurons and astrocytes in HCSs. Furthermore, our system demonstrates enhanced physiological relevance, as co-cultured brain organoids exhibit a higher level of the 4R-tau to 3R-tau ratio, closer to that found in adult human brain tissue. This addresses a key limitation of traditional brain organoid models, which typically retain fetal-like characteristics^28^. The gut’s influence on brain physiology was also notably evident at the transcriptomic level. When brain organoids were co-cultured with gut organoids, we observed a suppression of oxidative stress pathways, which aligns with growing recognition of the gut’s role in modulating CNS redox balance^71^. Additionally, there was an activation of neuronal activity pathways.

Metabolomic analysis revealed that the gut had elevated levels of amino acid metabolites. This elevation may explain the observed activation of neurons, as amino acids are important precursors of neurotransmitters and energy sources for neurons^72^. Furthermore, the role of the BBB as a metabolic filter was demonstrated, as it effectively attenuated the metabolic effects derived from the gut, highlighting the importance of including the BBB when modeling interactions between the peripheral and CNS^22^.

Additionally, our gut-BBB-brain co-culture model also proves to be a valuable research tool for AD studies. Our system successfully recapitulates key disease hallmarks that are not present in monoculture systems, including elevated Aβ and phosphorylated tau levels, altered tau isoform ratios, and transcriptomic profiles that align with clinical AD characteristics when compared to healthy controls. Notably, our system demonstrates the ability to detect changes in cognition-related pathways associated with the disease, representing a significant advancement over traditional models that often overlook these clinically relevant phenotypes^73^. These findings underscore the importance of considering peripheral organ interactions when investigating CNS disorders.

The most translational finding that arises from these experiments is that co-culture AD gut organoids with healthy brain organoids. This combination induced AD-like pathology in the brain organoids, providing experimental support for the emerging “gut-first” hypothesis in the AD field. This hypothesis suggests that gut-derived factors may initiate or worsen brain pathology^74^.

Our research addresses two significant gaps in neurodegeneration research: the absence of human-relevant multi-organ models and the need to understand how peripheral factors contribute to CNS pathology. By demonstrating that gut-brain interactions can influence AD pathologies, we provide experimental evidence supporting a link between gut health and AD risk.

While this represents a significant advancement, limitations still reflect broader challenges in the field. The absence of microbiota restricts the investigation of microbiome-gut-brain interactions, which are crucial to understanding the pathophysiology of AD^75^. To scale up the system for drug screening applications, it will be necessary to address the current time-intensive protocols, a challenge that many organoid-based platforms face^15^. We also face key challenges such as standardization, as organoid variability remains an issue, and scaling for high-throughput applications. However, integrating single-cell omics and AI-assisted analysis is rapidly addressing these hurdles^76^. Meanwhile, further verification of other cell lines will help extend our biological findings.

This study establishes a novel paradigm for investigating neurodegenerative diseases through the lens of inter-organ communication. By demonstrating that gut-derived factors can induce AD-relevant changes in brain tissue, we provide compelling evidence for viewing AD as a systemic disorder rather than just a condition of the CNS. Our microphysiological platform opens new opportunities to dissect gut-brain signaling mechanisms, validate clinical observations, and develop therapies that target the gut-brain axis — an approach that could transform our understanding and treatment of neurodegenerative diseases.

## 4. Materials and Methods

### 4.1 Human cell lines and differentiation

iPSC lines derived from healthy controls were obtained from Coriell, and iPSC lines derived from AD patients were kind gifts from Jian Zhao’s lab (Shanghai Institute for Advanced Immunochemical Studies, ShanghaiTech University) ^53–55^. All human iPSCs were cultured in feeder-free conditions in StemFlex medium (Thermo-Fisher, A3349401). iPSCs were passaged at 60-80% confluence using Gentle Cell Dissociation Reagent (STEMCELL Technologies, 07174) for 3 min and reseeded 1:10 onto Matrigel-coated (BD Biosciences, 354277) plates.

#### 4.1.1 Brain organoid differentiation

Differentiation of iPSCs into HCSs was performed according to the published protocol^27^. iPSCs were dissociated with Accutase (STEMCELL Technologies, 07920) into single cells and resuspended with StemFlex medium, adding 10 μM Y-27632 (Selleck Chemicals, S1049). 3.0 x 10^6^ cells were added into a single well of AggreWell^TM^800 plate (STEMCELL Technologies, 34815), corresponding to 1.0 x 10^4^ per microwell in StemFlex medium with 10 μM Y-27632. The plate was then centrifuged at 300 x g for 5 min and incubated overnight. The next day, aggregated cell spheroids were transferred into non-treated 100 mm plastic plates (JETBIOFIL, TCD000100) in StemFlex medium for 48 h. For neural induction, StemFlex medium was replaced with hPS medium (Advanced DMEM/F12 (Gibco, 11330057), 20% knockout serum (Gibco, 10828-028), 1x GlutaMAX (Gibco, 35050-061), 1x non-essential amino acids (Gibco, 11140050), 1x penicillin/streptomycin (Life Technologies, 15140122), and 100 μM 2-mercaptoethanol (Sigma-Aldrich, M3148)) supplemented with 5 μM Dorsomorphin (Selleck Chemicals, S7840) and 10 μM SB-431532 (Selleck Chemicals, S1067) on day 0. No medium changes were performed on day 1, and medium changes were performed daily for the next four days. On day 6, culture medium was replaced with neural medium (neurobasal medium (Gibco, 21103-049), 1x B27 supplement minus vitamin A (Gibco, 12587001), and 1x penicillin/streptomycin) supplemented with 20 ng/ml FGF2 (NovoProtein, C046) and 20 ng/ml EGF (NovoProtein, C029) for the next 19 days with daily changes for the first 10 days and every other day changes for subsequent 9 days. 2. To promote neural differentiation, FGF2 and EGF were replaced with 20 ng/ml BDNF (NovoProtein, C076) and 20 ng/ml NT3 (NovoProtein, C079) starting on day 25 and changed medium every other day. From day 43 onwards, HCSs were maintained in unsupplemented neural medium and with medium changed every 4-6 days. HCSs cultured for more than 100 days were used in subsequent experiments.

#### 4.1.2 Gut organoid differentiation

Differentiation of iPSCs into HIOs was carried out as previously described^77,78^. Briefly, iPSCs were cultured with DE medium consisting of RPMI1640 (Invitrogen, C11875500BT) containing 1x non-essential amino acids, 2% FBS (Sigma-Aldrich, F0193), and 100 ng/ml Activin A (NovoProtein, C687) for 3 consecutive days when iPSCs were 85%-90% confluent and were largely differentiation free. Starting on day 4, cells were fed daily with MHE medium consisting of RPMI 1640 medium containing 1x non-essential amino acids, 2% FBS, 500 ng/ml FGF4 (NovoProtein, CR08), and 3 μM CHIR99021 (Selleck Chemicals, S1263) for 4 days to generate mid-/hind-gut endoderm. On day 7, the cell pellet was dissociated with Accutase into single cells and resuspended with HIO medium (Advanced DMEM/F12, 1x B27 supplement (Gibco, 17504001), 1x N2 supplement, 1x penicillin/streptomycin, 1x GlutaMAX, adding 100 ng/ml Noggin, 100 ng/ml EGF, and 500 ng/ml R-Spondin1) with 10 μM Y-27632. For aggregation, 9.0 x 10^5^ cells were seeded into a well of AggreWell^TM^800 plate, corresponding to 3,000 cells per microwell in HIO medium with 10 μM Y-27632. The plate was then centrifuged at 300 x g for 5 min and incubated overnight. On day 8, aggregated cell spheroids were collected and embedded in 50 μL Matrigel (ABWBIO, 082755). Matrigel droplets were then plated in a single well of a 24-well dish, where HIO medium was used and replaced every 2-3 days. HIOs were manually passaged to reduce tissue density approximately every 14 days. HIOs cultured for more than 35 days were used in subsequent experiments.

#### 4.1.3 Astrocyte differentiation

Astrocyte differentiation was adapted from Blanchard et al^79^. First, for NPCs induction, StemFlex medium was changed to NPC medium (Advanced DMEM/F12: Neurobasal = 1:1, 1x N2 supplement (Gibco, 17502001), 1x B27 supplement minus vitamin A, 100 μM Ascorbic acid (Sigma, A4544), 1x GlutaMAX, 1x penicillin/streptomycin, and 100 μM 2-mercaptoethanol) supplemented with 2 μM SB431542, 3 μM CHIR99021, and 0.2 μM LDN193189 (Selleck Chemicals, S7507) for 6 days when iPSCs were 30%-45% confluent and medium was changed every other day. NPCs were then differentiated into astrocytes by seeding dissociated single cells at a 1.5 x 10^4^ cells/cm^2^ density onto Matrigel-coated plates in astrocyte medium (ScienCell, 1801). Astrocyte medium was changed every other day.

#### 4.1.4 BMEC differentiation

BMEC differentiation referred to Blanchard et al^79^. iPSCs were dissociated with Accutase into single cells and reseeded at 3.5 × 10^4^ cells/cm^2^ onto Matrigel-coated plates in StemFlex medium with 10 μM Y-27632. For the next 2 days, medium was replaced with StemFlex medium daily. On day 3, the medium was changed to DeSR1 medium (Advanced DMEM/F12, 1x GlutaMAX, 100 μM 2-mercaptoethanol, 1x non-essential amino acids, 1x penicillin-streptomycin, and 6 μM CHIR99021). For the following 5 days, the medium was changed to DeSR2 medium (Advanced DMEM/F12, 1x GlutaMAX, 100 μM 2-mercaptoethanol, 1x non-essential amino acids, 1x penicillin-streptomycin, and 1x B27 supplementary) and changed every day. After 5 days of DeSR2, the medium was changed to hECSR1 (Human Endothelial SFM (Thermo Scientific, 11111044) supplemented with 1x B-27 supplement, 10 μM retinoic acid (Sigma, R2625), and 20 ng/ml FGF2).

#### 4.1.5 Pericyte differentiation

Pericyte differentiation also referred to Blanchard et al^79^. iPSCs were dissociated with Accutase into single cells and reseeded at 4.0 × 10^4^ cells/cm^2^ onto Matrigel-coated plates in StemFlex medium with 10 μM Y-27632. For the next 3 days, medium was changed to PC medium (Advanced DMEM/F12: Neurobasal = 1:1, 1x N2 supplement, 1x B27 supplement minus vitamin A, and 1x penicillin/streptomycin) supplemented with 25 ng/ml BMP4 (NovoProtein, CR93) and 8 μM CHIR99021. On days 4 and 5, medium was changed to PC medium supplemented with 10 ng/ml PDGF-BB (NovoProtein, C199) and 2 ng/ml Activin A.

### 4.2 iBBB construction and permeability studies

2x 10^5^ astrocytes and 2x 10^5^ pericytes were mixed together and encapsulated in 50 μL BBB Matrigel (Matrigel: Advanced DMEM/F12 = 1: 2). Matrigel cell solution was then seeded onto the basal side of 12-well trans-well polyester membrane cell culture inserts (0.4 μm pore size, LABSELECT, 14212) and allowed to solidify for 40 min at 37 ℃ and then grow in neural medium. The next day, 1.13 x 10^6^ BMECs were seeded onto the apical side of the Matrigel-coated trans-well and cultured in hECSR1.

#### 4.2.1 TEER measurements

TEER value was measured daily using an EVOM2 volt/ohmmeter with STX2 electrode set (World Precision Instruments, USA). The iBBB models with barrier function less than 1000 Ω·cm^2^ were excluded from subsequent tests and co-culture.

#### 4.2.2 Trans-BBB flux of FITC-dextran

1mg/ml FD-4 or FD-70 was added to the apical side of iBBB, and medium from the basal side was withdrawn after 1, 2, and 3 hours. Fluorescence intensity was measured by a multi-mode microplate reader (λex 488 nm, λem 515 nm). Determine the amount of permeated FD molecules by comparison of the observed fluorescence values with a calibration curve produced with known concentrations. Calculate the apparent permeability coefficient (Papp) from the mean flux values according to:

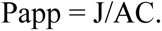

where J is the flux of the molecule (moles/sec), A is the permeation area (cm^2^), and C is the concentration of the molecule in the upper compartment (moles/cm^3^).

#### 4.2.3 Efflux pump activity

MK-571 and PSC-833 were used as inhibitors of MRPs and P-gp, respectively, whereas H2DCFDA and Rhodamine123 are substrates of MRPs and P-gp, respectively. iBBB was incubated with 10 μM MK-571 or 10 μM PSC-833 on the apical side at 37 ℃ for 1 hour, and then 10 μM H2DCFDA or 10 μM Rhodamine123 was added to the apical side for another hour. Remove 200 μL from the basal side and measure fluorescence intensity by a multi-mode microplate reader (λex 492 nm, λem 527 nm for H2DCFDA; λex 488 nm, λem 540 nm for Rhodamine123).

#### 4.2.4 Permeability Assays

40 μg/ml human albumin (NovoProtein, CP01), 10 μg/ml IgG (NovoProtein, CX82), 2.5 μg/ml transferrin (NovoProtein, CJ41), or 1 μg/ml Aβ (MedChem Express, HY-P1363) were added into the apical side of iBBB at 37 ℃ overnight. The next day, samples from the apical side and the basal side were collected, respectively, and quantification of the proteins was performed using the following ELISA kits from JONLNBIO: albumin (JL15013), IgG (JL12687), transferrin (JL18359), and Aβ (JL14446).

#### 4.2.5 Drug permeability tests

The TEER was measured before carrying out permeability tests. Cells were incubated with Hank’s Balanced Salt Solution (HBSS) (Thermo Scientific, 14175079) for 2 hours. Drugs at a concentration of 10 μM in Table S2 diluted in HBSS were added to the apical side of the iBBB. After incubation at 37 ℃ for 60 min, samples from the apical side and the basal side were collected, respectively. Drug concentrations were measured by LC/MS analysis.

### 4.3 Common culture medium

The gut compartment of the GBB microphysiological system was fed with HIO medium, and the brain compartment was fed with neural medium.

### 4.4 Cryopreservation

Organoids were fixed in 4% paraformaldehyde (PFA) (Solarbio, P-1110) for 1 hour. They were then washed with PBS and transferred into a 30% sucrose solution and kept at 4 ℃ for 48-72 hours. Subsequently, they were transferred into Tissue Freezing Medium (Tissue Plus O.C.T. Compound, scigen, 4586) and snap-frozen and stored at −80 ℃. For immunofluorescence, 16 μm-thick sections were obtained using a cryostat.

### 4.5 Immunofluorescence staining

Cells were fixed in 4% PFA for 15 min and then washed with PBS. Cryosections were washed with PBS to remove excess tissue freezing medium. Cells and cryosections were then blocked in 10% normal goat serum (NGS), 0.3% Triton X-100 diluted in PBS for 1 hour at room temperature. Next, they were incubated with primary antibodies diluted in PBS containing 5% NGS and 0.1% Triton X-100 at 4 ℃ overnight. They were incubated with secondary antibodies diluted in PBS containing 5% NGS and 0.1% Triton X-100 at room temperature for 1 h after they were washed with PBS. Cells were then counterstained with nuclear dye DAPI (1:1000, Thermo-Fisher, D1306) for 5 min. The antibodies used in this study are shown in Table S3. Confocal fluorescent images were acquired using a Leica Confocal microscope and an Olympus SpinSR spinning disk. Image analysis was performed using ImageJ software.

### 4.6 Western blot

Cells or tissues were lysed in RIPA (Beyotime, P0013B) supplemented with protease inhibitors (Beyotime, ST506). Protein concentration was measured using the BCA assay (Thermo Fisher Scientific, 23227). Adjust samples to equal concentrations with lysis buffer and then mix samples with 5x Omni-Easy™ Protein Sample Loading Buffer (Epizyme, LT101). Samples were loaded into 10-12% SDS-polyacrylamide gels and then transferred onto PVDF membranes (Millipore, IPVH00010). Membranes with proteins were incubated in 10% non-fat milk in TBST for 30 min at room temperature. Next, they were incubated with primary antibodies diluted in TBST containing 5% milk at 4 ℃ overnight. Primary antibody used was mouse anti-Phospho-Tau (ThermoFisher, MN1020; 1:1000). The membranes were incubated with secondary antibodies diluted in TBST containing 5% milk at 4 ℃ overnight after they were washed with TBST. Apply enhanced chemiluminescence (ECL) substrate (Sharebio, SB-WB001) evenly onto the membrane and capture signals using a chemiluminescence imaging system.

### 4.7 Semi-quantitative PCR

cDNA was generated from RNA and amplified using primers^80^ flanking exon 10 (forward 5’-AAGTCGCCGTCTTCCGCCAAG-3’; reverse 5’-GTCCAGGGACCCAATCTTCGA-3’). The PCR product was then run on a 2% agarose gel with 381bp and 288bp fragments, indicating 4R and 3R, respectively. Gray values were then calculated using ImageJ.

### 4.8 RNA-sequencing

RNA-sequencing was performed by Azenta Life Science (Shanghai, China). 1 μg total RNA was used for the following library preparation. The poly(A) mRNA isolation was performed using Oligo(dT) beads. The mRNA fragmentation was performed using divalent cations and high temperature. Priming was performed using Random Primers. First-strand cDNA and second-strand cDNA were synthesized. The purified double-stranded cDNA was then treated to repair both ends and add a dA-tailing in one reaction, followed by a T-A ligation to add adaptors to both ends. Size selection of Adaptor-ligated DNA was then performed using DNA Clean Beads. Each sample was then amplified by PCR using P5 and P7 primers, and the PCR products were validated. Then libraries with different indexes were multiplexed and loaded on an Illumina HiSeq instrument for sequencing using a 2x150 paired-end (PE) configuration according to the manufacturer’s instructions. Differential expression analysis used the DESeq2 Bioconductor package, a model based on the negative binomial distribution. The estimates of dispersion and logarithmic fold changes incorporate data-driven prior distributions; p.adj of genes were set ≤ 0.05 to detect differentially expressed ones. GOSeq was used to identify Gene Ontology (GO) terms that annotate a list of enriched genes with a significant p.adj less than or equal to 0.05. KEGG (Kyoto Encyclopedia of Genes and Genomes) is a collection of databases dealing with genomes, biological pathways, diseases, drugs, and chemical substances (http://en.wikipedia.org/wiki/KEGG). We used scripts in-house to enrich significantly differentially expressed genes in KEGG pathways.

### 4.9 Metabolomics

Non-targeted metabolomic analysis and bioinformatic data processing were performed by Tsingke (Beijing, China).

#### 4.9.1 Sample preparation

The sample was thawed on ice. A 500 μL solution (Methanol: Water = 4:1, V/V) containing internal standard was added to the sample and vortexed for 3 min. The sample was placed in liquid nitrogen for 5 min and on the dry ice for 5 min, and then thawed on ice and vortexed for 2 min. This freeze-thaw cycle was repeated three times in total. The sample was centrifuged at 12000 rpm for 10 min at 4 ℃. A 450 μL of the supernatant was transferred and concentrated. A 100 μL solution (Methanol: Water = 7:3, V/V) was used to reconstitute the sample. Then the sample was vortexed for 3 minutes, and sonicated for 10 minutes in an ice bath. The sample was then centrifuged at 12000 rpm.

#### 4.9.2 HPLC conditions

All samples were for two LC/MS methods. One aliquot was analyzed using positive ion conditions and was eluted from T3 column (Waters ACQUITY Premier HSS T3 Column 1.8µm, 2.1 mm x 100 mm) using 0.1% formic acid in water as solvent A and 0.1 % formic acid in acetonitrile as solvent B in the following gradient: 5 to 20 % in 2 min, increased to 60% in the following 3 mins, increased to 99 % in 1 min and held for 1.5 min, then come back to 5% mobile phase B within 0.1 min, held for 2.4 min. The analytical conditions were as follows: column temperature, 40 ℃; flow rate, 0.4 mL/min; injection volume, 4 μL. Another aliquot was using negative ion conditions and was the same as the elution gradient of positive mode.

#### 4.9.3 MS conditions

All the methods alternated between full scan MS and data-dependent MS scans using dynamic exclusion. MS analyses were carried out using electrospray ionization in the positive ion mode and negative ion mode using full scan analysis over m/z 75-1000 at 35000 resolutions.

### 4.10 Statistical analysis

All data are presented as mean ± SEM. P values were calculated by two-tailed Student’s t-tests. Significance is expressed as *P<0.05, **P<0.01, ***<0.001, ****P<0.0001.

## Acknowledgements

We thank the staff members of the Integrated Laser Microscopy System (https://cstr.cn/31129.02.NFPS.CLMIS) at the National Facility for Protein Science in Shanghai (https://cstr.cn/31129.02.NFPS) for providing technical support and assistance in data collection and analysis. We acknowledge the iPSCs provided by Dr. Jian Zhao’s laboratory of ShanghaiTech. We thank Dr. Li Tan’s laboratory of Chinese Academy of Sciences for the technical support on LC-MS provided by Dr. Huai-Jiang Xiang.

## Data Availability Statement

The datasets generated and analyzed during the current study are available in the GEO repository, GSE311936.

## Funding

The National Key Research and Development Program of China (2024YFA1108000) The National Natural Science Foundation of China (82441053)

The Shanghai Key Laboratory of Aging Studies (19DZ2260400)

Shanghai Municipal Science and Technology Major Project (2019SHZDZX02)

